# Optogenetics reprogramming of planktonic cells for biofilm formation

**DOI:** 10.1101/229229

**Authors:** Aiguo Xia, Shuai Yang, Yajia Huang, Zhenyu Jin, Lei Ni, Lu Pu, Guang Yang, Fan Jin

## Abstract

Single-cell behaviors play essential roles during early-stage biofilms formation. In this study, we evaluated whether biofilm formation could be guided by precisely manipulating single cells behaviors. Thus, we established an illumination method to precisely manipulate the type IV pili (TFP) mediated motility and microcolony formation of *Pseudomonas aeruginosa* by using a combination of a high-throughput bacterial tracking algorithm, optogenetic manipulation and adaptive microscopy. We termed this method as Adaptive Tracking Illumination (ATI). We reported that ATI enables the precise manipulation of TFP mediated motility and microcolony formation during biofilm formation by manipulating bis-(3′-5′)-cyclic dimeric guanosine monophosphate (c-di-GMP) levels in single cells. Moreover, we showed that the spatial organization of single cells in mature biofilms can be controlled using ATI. Thus, the established method (i.e., ATI) can markedly promote ongoing studies of biofilms.

## Introduction

Biofilm is the most successful lifestyle that allows microbes to survive and thrive in nature (H. C. Flemming et al., 2016; Hall-Stoodley, Costerton, & Stoodley, 2004; Hibbing, Fuqua, Parsek, & Peterson, 2010). Biofilm formation has been reported to occur in several stages; namely, the transition from free-living single cells to complex microcolonies, secretion of extracellular polymeric substance (EPS) to enclose microcolonies (H.-C. Flemming & Wingender, 2010), differentiation and development of three-dimensional morphology (Stewart & Franklin, 2008), and, ultimately, the release of dispersal cells to restart the lifecycle (McDougald, Rice, Barraud, Steinberg, & Kjelleberg, 2012). Recent studies have indicated that single-cell motility mechanisms (Gibiansky et al., 2010), including type IV pili (TFP)- or flagella-mediated motility (Conrad et al., 2011), and EPS production (Zhao et al., 2013) play essential roles in determining the location and time of microcolony formation. These fundamental findings not only provide researchers with an excellent opportunity for developing novel strategies to prevent biofilm formation on medical and industrial settings, which can cause antibiotic-tolerant infections (Costerton, Stewart, & Greenberg, 1999) and the destruction of flow systems, but also motivate researchers to determine whether biofilm formation can be completely controlled by precisely manipulating single-cell motility and microcolony formation during early-stage biofilm formation.

Leifer et al. reported that a combination of optogenetic manipulation (Leifer, Fang-Yen, Gershow, Alkema, & Samuel, 2011) and adaptive microscopy (Tischer, Hilsenstein, Hanson, & Pepperkok, 2014) enables the direct manipulation of free-moving nematode *Caenorhabditis elegans*, thus motivating the development of a method that can directly manipulate many moving bacteria on a surface during biofilm formation. Similarly, in this study, using a combination of a high-throughput bacterial tracking algorithm, optogenetic manipulation, and adaptive microscopy, we first established a method, adaptive tracking illumination (ATI). This method could precisely illuminate single moving cells through the *in-situ* analysis and tracking of moving bacterial trajectories. Subsequently, we applied this method to directly manipulate the TFP-mediated motility and microcolony formation of single *Pseudomonas aeruginosa* cells during early-stage biofilm formation. Bis-(3′-5′)-cyclic dimeric guanosine monophosphate (c-di-GMP) signaling pathways (Hengge, 2009) are established to govern the pleiotropic behavior of bacteria, including bacterial motility, EPS production, and biofilm formation. Therefore, we modified the genome of *P. aeruginosa* by using an optogenetic tool (M. H. Ryu & Gomelsky, 2014), which facilitated the direct manipulation of the c-di-GMP level in single cells by using near-infrared light. We observed that precisely manipulating the c-di-GMP level in single cells during early-stage biofilm formation enabled the guiding of subsequent organization of cells in mature biofilms. Thus, the established method (i.e., ATI) can markedly promote modern research on biofilms, ranging from fundamental to industrial studies.

## Results

### ATI enables precise illumination of single *P. aeruginosa* cells

During biofilm formation, *P. aeruginosa* cells differently deploy their TFP to mediate distinct motility, with a mobile or an immobile phenotype (Ni et al., 2016). The process involves the frequent dispersal of mobile cells on surfaces, enabling them to find a habitat that can supply sufficient nutrients. By contrast, immobile cells stall in their place to form microcolonies. These distinct TFP-mediated motility phenotypes are thereby expected to affect subsequent biofilm formation, including the organization and differentiation of cells and the structure of mature biofilms. To precisely illuminate single *P. aeruginosa* cells with distinct motility, we first acquired the bright-field images of the cells and then tracked them to obtain their trajectories in real time. This real-time information was further analyzed to identify mobile and immobile cells. Thereafter, information on the regions containing selected cells with the motility phenotype of interest was input into a digital micromirror device to generate a mask. The mask was finally projected on these selected cells through an additional objective. We termed this illumination as ATI (Figure 1a, Figure 1—figure supplement 1 and Figure 1—figure supplement 2). Our results indicated that feedback illuminations could generate projected patterns to exactly follow the cell movement (Figure 1b and Video 1) or single cells divisions (Figure 1c and Video 2) in real time.

**Figure 1 with 2 supplements.**
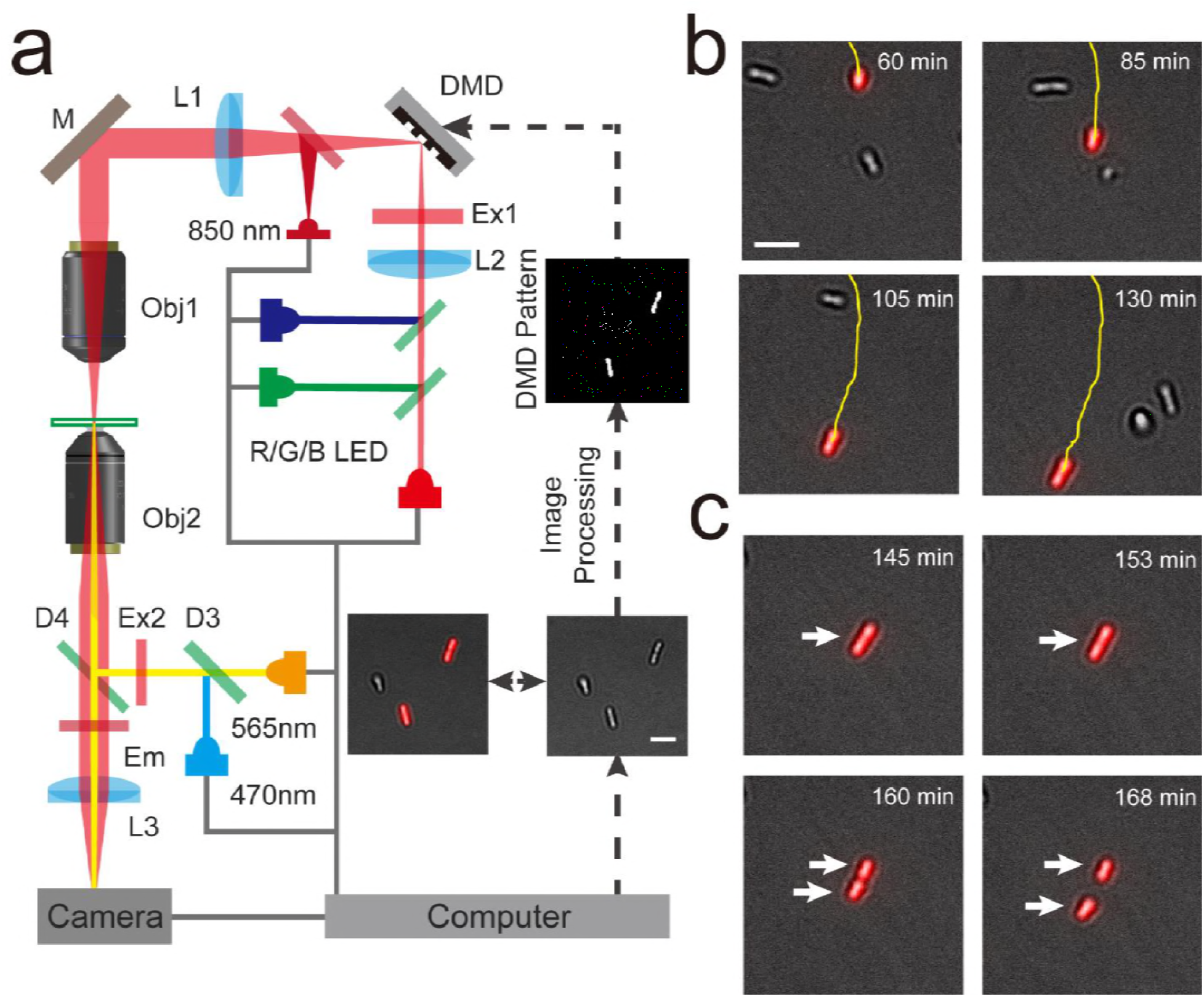
Using Adaptive Tracking Illuminations (ATI) to exactly illuminate single *P. aeruginosa* cells on surface. **(a)** Schematic drawing of the ATI system. A high-throughput bacterial tracking algorithm was employed for analyzing cells’ behaviors in real time and the information were immediately feedback to an adaptive microscope equipped with a digital micromirror device (DMD). **(b)** One interested cell being tracked and projected in real time is depicted. **(c)** The feedback illuminations also can generate projected patterns to exactly follow the daughter cells after the division of one interested cell. Scale bars for all images are 5μm.

### ATI enables precise manipulation of TFP-meditated motility and microcolony formation of single cells

To directly manipulate the TFP-meditated motility of the selected single cells, we incorporated an optogenetic part into the chromosome of *P. aeruginosa* by using the mini-CTX system(Hoang, Kutchma, Becher, & Schweizer, 2000). This part encodes a heme oxygenase (*bphO*) and c-di-GMP diguanylate cyclase(*bphS*) that can cyclize two GTP molecules to form a c-di-GMP molecule in the presence of near-infrared light (M. H. Ryu & Gomelsky, 2014). Furthermore, the optogenetic part enabled the direct manipulation of intracellular c-di-GMP levels by using illumination at 640 nm (Figure 2a). Elevation of c-di-GMP levels is known to repress bacterial motility and promote biofilm formation (Hengge, 2009). To optimize the intensity of manipulation lights, we varied the LED light intensity (640 nm) to stimulate optogenetically modified bacteria. Furthermore, we monitored c-di-GMP levels in single cells by using a fluorescent reporter (Rybtke et al., 2012). The c-di-GMP reporter encodes two fluorescent proteins. A green fluorescent protein (GFP) is expressed using a c-di-GMP regulatory promoter (PcdrA), and a red fluorescent protein (mCherry) is expressed using a constitutive promoter (P*A1O4O3;* Figure 2a). Thus, the ratio of fluorescent intensities derived from GFP (F_G_) and mCherry (F_R_) was used to determine the c-di-GMP levels in single cells. We observed that c-di-GMP levels, as indicated by the Fg/Fr ratio, markedly increased 10 folds after 4 hours of illumination (> 0.02 mW.cm^−2^) with 640-nm LED lights (Figure 2— figure supplement 1a). Notably, the growth of cell was not affected even using the illumination at 0.26 mW.cm^−2^ (Figure 2—figure supplement 1b).

**Figure 2 with 2 supplements.**
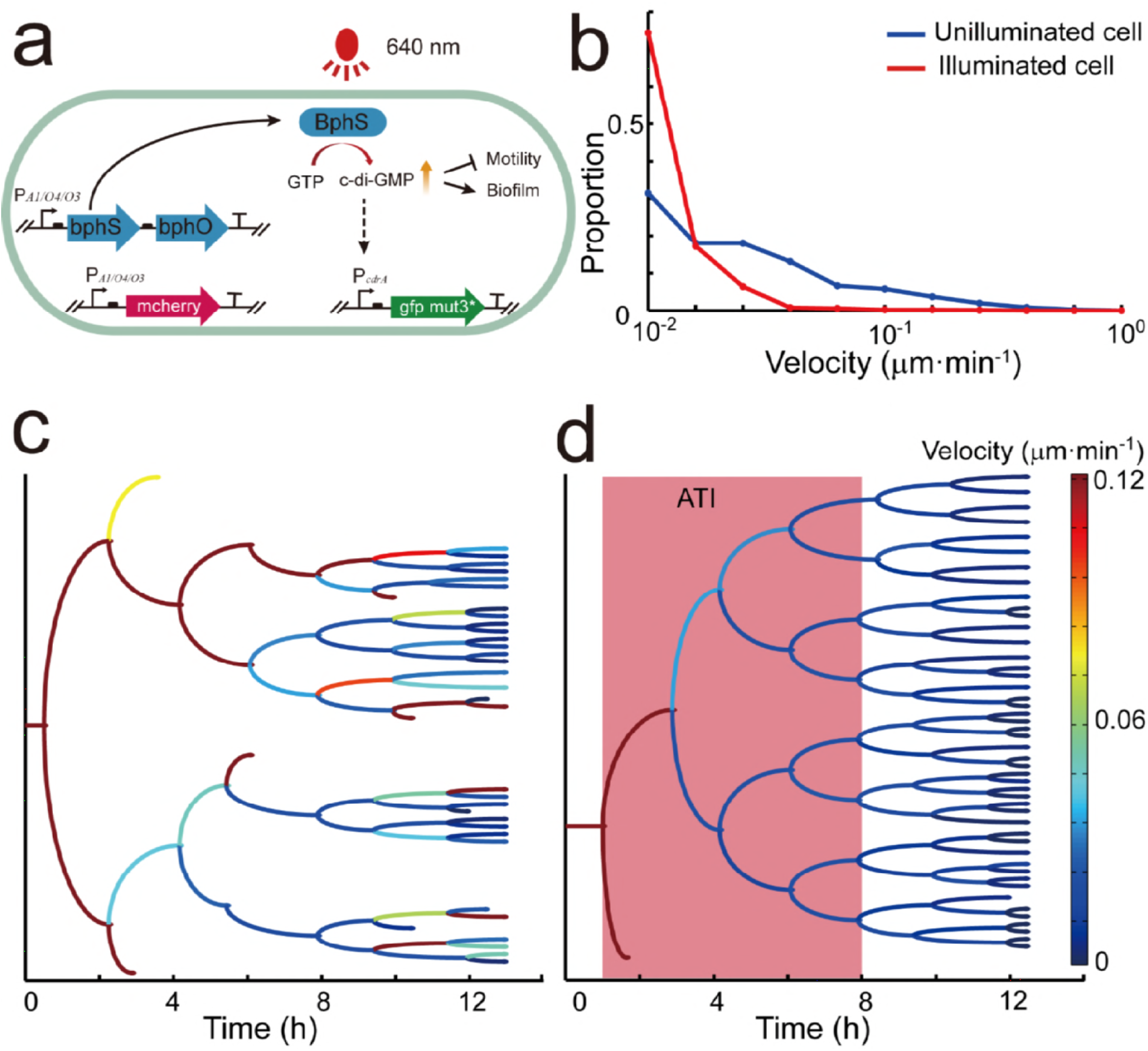
By introducing optogenetic modules, the ATI can precisely manipulate TFP meditated motility. **(a)** Schematic of optogenetic circuit. Optogenetic modules encode a c-di-GMP diguanylate cyclase (BphS), which allowed us to manipulate c-di-GMP level using an illumination of 640 nm lights. Expression of GFP mut3* under control of the c-di-GMP regulatory promoter (PcdrA) was used for monitoring the c-di-GMP levels. A mCherry reporter fused to the constitutive promoter P*A1/O4/O3* served as an internal control. **(b)** The illuminated cells have much slower velocity than unilluminated cells. Data were analyzed and obtained after 4 hours of manipulation by ATI. **(c)** Genealogical tree of one unilluminated cell and **(d)** one illuminated cell. The velocity of each cell is color-coded along each lineage. ATI enabled mobile cells and their offspring to switch to immobile phenotype as indicted by their averaged moving velocities decreased from 0.2 to 0.01 μm·min^−1^ and less detachment events. Experiments in **b-d** were carried out at least three times and one representative example is shown.

We first selected 33–66% of single cells with mobile motility and illuminated them (0.05 mW.cm^−2^) along with their offspring for 7 hours by using ATI. Treatment with ATI for 3 hours enabled all mobile cells and their offspring to switch to the immobile phenotype, as indicated by their average moving velocities, which decreased from 0.2 to 0.01 μm·min^−1^ (Figs. 2b and d and Video 3). This observation contrasted with the observation that 50% of mobile cells (> 0.03 μm·min^−1^) without ATI treatment remained mobile on surfaces (Figure 2b and c and Video 3). Wild-type *P. aeruginosa* can spontaneously switch between mobile and immobile phenotypes during early-stage biofilm formation(Mikkelsen, Sivaneson, & Filloux, 2011), which results in that the velocity distribution arising from single cells are broadly distributed from 0.01 to 0.5 μm·min^−1^ (Figure 2b). Moreover, we observed that manipulation by using ATI markedly enhanced the possibility that the post-division cells remained attached to surfaces (Figure 2d). This observation contrasted with the observation that unilluminated cells exhibited asymmetrical detaching behaviors after division (Figure 2c): one daughter cell preferred to remain attached to surfaces, whereas another cell preferred to detach from surfaces.

Simultaneously, we observed that 7 hours of ATI treatment considerably increased the c-di-GMP levels in these illuminated cells and their offspring, as indicated by the F_G_/F_R_ ratio, which increased from 1.5 to 7.5 (Figure 3a, d and Video 3). This observation was in sharp contrast to the observation that c-di-GMP levels remained low in unilluminated mobile cells (Figure 3a, c and Video 3). The mCherry-induced fluorescence remained nearly constant in the illuminated and unilluminated cells (Figure 2—figure supplement 2). Notably, the optogenetic manipulation of single mobile cells and their offspring enabled these cells to stall in their place to form a microcolony during biofilm formation, whereas mobile cells and their offspring without optogenetic manipulation were more likely to move and spread on surfaces (Figure 3b and Video 3). To evaluate how optogenetic manipulation affects microcolony formation, we calculated the clustering ability of illuminated and unilluminated cells through hierarchical clustering. Clustering calculation indicated that ATI enabled single mobile cells to form a microcolony, as indicated by the distance among the daughter cells (< 4 μm; Figure 3d and f). This observation was in sharp contrast to the observation that unilluminated cells and their offspring were more likely to spread on surfaces, as indicated by the distance among these daughter cells (> 40 μm; Figure 3c and e).

**Figure 3.**
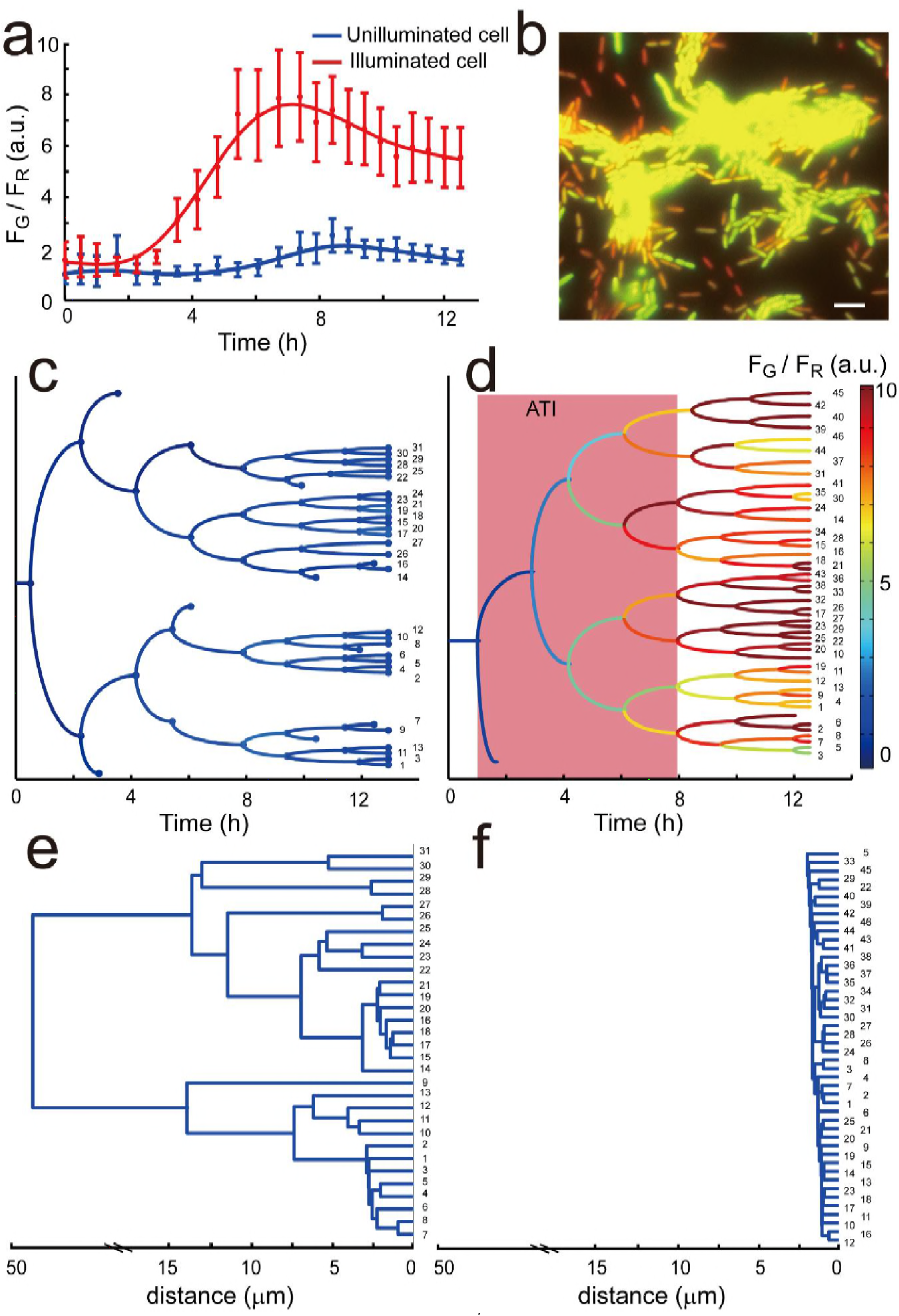
c-di-GMP level was manipulated using ATI system in single cells. **(a)** The illuminated cells and their offspring have fivefold higher levels of c-di-GMP than unilluminated cells after 6 hours as indicated by the contrast of Fg/Fr. Error bars represent means ± s. d with n = 3 biological replicates. **(b)** A merged image of mCherry and GFP mut3* fluorescence microscopy image was presented after 14 hours. The illuminated cells have brighter green fluorescence, which indicates higher c-di-GMP level. Scale bar were set at 5μm. The c-di-GMP level from one mother cell is depicted by Genealogical trees for unilluminated **(c)** and illuminated cell **(d)**, respectively. **(e)** and **(f)** for the corresponding hierarchical clustering analysis in **(c)** and **(d)**. After manipulation by ATI for 12 hours, the distance among the daughter cells was less than 4 μm, which is in clear contrast to the largest distance between unilluminated daughter cells larger than 40 μm. Experiments in **b-f** were carried out at least three times and one representative sample is shown.

### ATI enables the guiding of biofilm formation in *P. aeruginosa*

After 7 hours of illumination, c-di-GMP levels remained high in the cells and their offspring, even without further optogenetic manipulation (Video 3). This behavior guided subsequent biofilm organization by manipulating microcolony formation during early-stage biofilm formation. To demonstrate this phenomenon, we labeled the optogenetically modified cells by using a green or red fluorescent protein (EGFP or mCherry), which did not contain the fluorescent reporter of c-di-GMP. We cocultured these differently labeled cells in a flow cell to facilitate their biofilm formation, during which, we only selected GFP- or RFP-labeled cells and manipulated them by using ATI in the first 10 hours. Our results indicated that ATI enabled the selected cells (GFP- or RFP-labeled cells) and their offspring to form microcolonies in advance, whereas the unselected cells distributed around the formed microcolonies (Figure 4a). This result was in sharp contrast to that obtained for GFP- and RFP-labeled cells without ATI treatment (Figure 4—figure supplement 1). Thereafter, we continuously cultured these young biofilms (up to 72 hours) with a distinct organization of cells in dark to allow their maturation. These distinct young biofilms developed to mature biofilms possessing a distinct spatial organization of GFP- or RFP-labeled cells (Figure 4b). These results revealed that cell manipulation by using ATI during early-stage biofilm formation enabled the guiding of biofilm formation.

**Figure 4 with 1 supplements.**
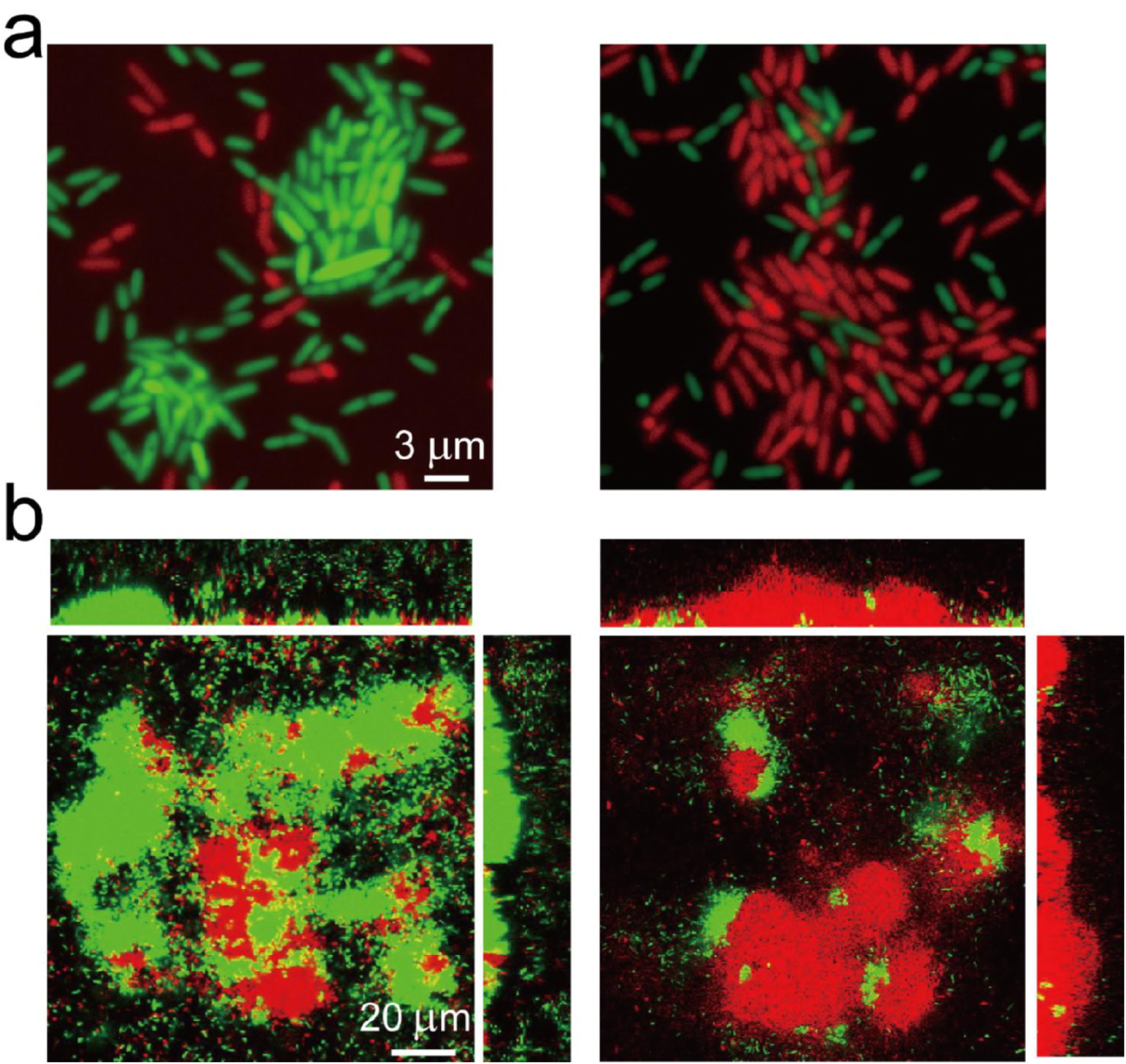
Guiding biofilm formation using ATI in *P. aeruginosa.* **(a)** ATI enables selected cells and their offspring to form microcolonies in advance. The optogenetic modified cells (do not contain fluorescent reporter of c-di-GMP) labeled by a green (EGFP) or red (mCherry) fluorescent protein were used for biofilm cultivation in flow cell. Left panel, EGFP-labeled cells were manipulated by ATI; Right panel, mCherry-labeled cells were selected for manipulation. The fluorescence image was attained in flow cell at the same time point about t~14 h. **(b)** The corresponding mature biofilm structure in **(a)** after 3 days culture. The selected GFP- or the RFP-labeled cells possess distinctive spatial organizations of biofilms. Experiments in **a-b** were carried out at least three times and one representative image is depicted.

## Discussion

We developed the ATI method that could be used to precisely manipulate TFP-mediated motility of single *P. aeruginosa* cells during early-stage biofilm formation. In this method, the bacteria were modified using an optogenetic part that enabled illumination with near-infrared light to directly regulate intracellular c-di-GMP levels. We showed that ATI could manipulate single cells with a mobile phenotype to switch to an immobile phenotype. Consequently, these manipulated cells could stall in their place to form microcolonies in advance, whereas unmanipulated cells with a mobile phenotype were more likely to move and spread on surfaces, facilitating the control of the location and time of early-stage biofilm formation. Accordingly, we showed that the spatial organization of single cells could be precisely controlled in a young biofilm of *P. aeruginosa* cells. Notably, our results indicated that the organization of single cells in young biofilms affected subsequent cell organization in mature biofilms, which enabled the further control of the structure and spatial organization of cells in biofilms. It should be emphasized that the strategy used herein to control the spatial organization of bacterial colonies is different than that using micro-threedimensional printing (Connell, Kim, Shear, Bard, & Whiteley, 2014).

In addition to using ATI to guide biofilm formation, we expect that our method can be used to answer various questions or resolve problems in microbiology. This is because: 1) The hardware used to build ATI, mainly an air objective, a commercial projector, and an LED controller, are quite common and inexpensive; 2) The optical setup of ATI is compatible with that of other commonly used microscopic techniques, including fluorescence, confocal, and total internal reflection microscopy; 3) The wavelengths in ATI can be easily expanded to multiple colors to adapt to different optogenetic tools (Ohlendorf, Vidavski, Eldar, Moffat, & Moglich, 2012; Olson, Hartsough, Landry, Shroff, & Tabor, 2014; M.-H. Ryu, Fomicheva, Moskvin, & Gomelsky, 2017); and 4) The software and algorithms used in ATI, including image processing, single-cell tracking, and analysis of phenotypes, are quite flexible to modifications. These factors allow researchers to simply integrate the ATI setup to their microscope and quickly modify the algorithms to track single cells with phenotypes of their interest, thus markedly prompting studies of optogenetic tools. Notably, the time required for data processing limits the application of ATI for investigating a quickly evolving bio-systems or cellular processes. For example, data processing of a live image (processed in a commercial desktop equipped with an intel i7 CPU) in the present study typically took 3 seconds, limiting the use of ATI for manipulating rapidly swimming bacteria, whose velocity can typically reach tens of microns per second. In addition to the data processing speed, the accuracy of the tracking algorithm limits the application of ATI for investigating microbes with high cell densities. For example, the current ATI cannot be used to manipulate the phenotypes of single cells during middle-stage biofilm formation because the algorithm cannot accurately track single cells in a dense microcolony. Developing new tracking algorithms or using powerful computers can address these limitations, thus considerably expanding ATI applications.

## Materials and Methods

### Strains and Growth conditions

Bacterial strains and plasmids used in this study are listed in Table 1. Strains were grown on LB agar plates at 37°C for 24 hours. Monoclonal colonies were inoculated and cultured with a minimal medium (FAB) at 37°C with 30 mM glutamate as carbon source under an aerobic condition, in which the medium contains following compositions per liter: (NH_4_)_2_SO_4_, 2 g/L; Na_2_HPO_4_-12H_2_O, 12.02 g/L; KH_2_PO_4_, 3 g/L; NaCl, 3 g/L; MgCl_2_, 93 g/mL; CaCl_2_.2H_2_O, 14 g/mL; FeCl3, 1 uM; and trace metals solution(CaSO_4_-2H_2_O, 200 mg/L; MnSO_4_-7H_2_O, 200 mg/L; CuSO_4_.5H_2_O, 20 mg/L; ZnSO_4_-7H_2_O, 20 mg/L; CoSO_4_.7H_2_O, 10 mg/L; NaMoO_4_.H_2_O, 10 mg/L; H_3_BO_3_, 5 mg/L) 1 mL/L. The strains were harvested at OD_600_ = 2.1, and the bacterial cultures were further diluted (1:100) in fresh FAB medium to OD_600_ = 0.4~0.6 before used.

**Table 1.**
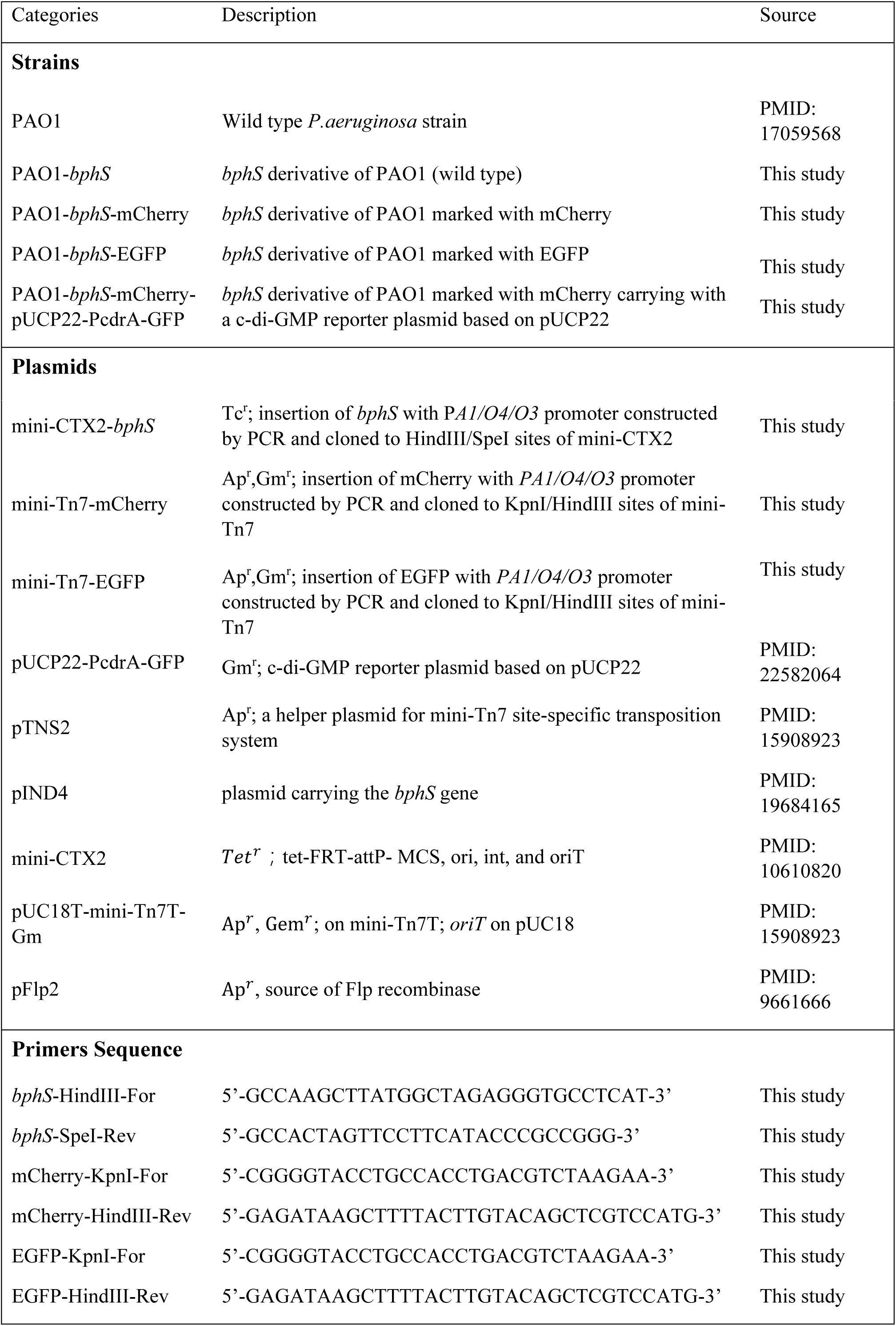
Strains, plasmids, and primers used in this study.

### Construction of optogenetics and c-di-GMP reporter strain in *P. aeruginosa*

Insertion mutant *bphS* was constructed by mini-CTX system using a modified procedure for *P.aeruginosa*. The *bphS* mutants marked with different fluorescent proteins were constructed by mini-Tn7 site-specific transposition system using a modified procedures for *P.aeruginosas*. We constructed unmarked insertion mutants by Flp-mediated excision of the antibiotic resistance marker. Firstly, *bphS* fragments obtained from the plasmid pIND4 was cloned into the vector mini- CTX2 with the *PA1/O4/O3* promoter in the upstream of MCS *via* a two-piece ligation. The constructed plasmid was electroporated into PAO1 and the corresponding recombinant strain was identified by screening on LB agar plates containing 1mM IPTG and 100 μg.mL^−1^ tetracycline. Thereafter, the strains were electroporated with a pFLP2 plasmid and distinguished on LB agar plates containing 5% (w/v) sucrose for the excision of resistance marker. Further, the *bphS* mutants were marked with mCherry/EGFP by using mini-Tn7 system as the similar procedure with mini-CTX. C-di-GMP reporter plasmid was electroporated to the mCherry/EGFP marked *bphS* mutants to determine the intercellular c-di-GMP level as required. The constructed plasmids and strains are listed in the Table 1.

### Setup of Adaptive Tracking Illuminations

We schematically show the setup of the Adaptive Tracking Illuminations (ATI) (Figure 1a). More specifically, an inverted fluorescent microscope (Olympus, IX71) was modified to build the ATI. The modification includes that: 1) A commercial DMD-based LED projector (Gimi Z3) was used to replace the original bright-field light source, in which the original lens in the projector were removed and the three-colors (RGB) LEDs were rewired to connect to an external LED driver (ThorLabs) controlled by a single chip microcomputer (Arduino UNO r3); 2) The original bright-field condenser was replaced with an air objective (40×, NA = 0.6, Leica); and 3) an additional 850 nm LED light (ThorLabs) was coupled to the illumination optical path using a dichroic mirror (Semrock) for the bright-field illuminations. Note that 850 nm LED light is a safe light to ensure that bright-field illuminations do not affect the optogenetics manipulation. The inverted fluorescent microscope equipped with a 100×oil objective and a sCMOS camera (Zyla 4.2 Andor) was used to collect bright-field images with 0.2 frame rate. The bright-field images were further analyzed to track multiple single cells in real time using a high-throughput bacterial tracking algorithm coded by Matlab. The projected contours of selected single cells were sent to the DMD (1280 × 760 pixels) that directly controlled by a commercial desktop through a VGA port. The manipulation lights were generated by the red-color LED (640 nm), and were projected on the single selected cells in real time through the DMD, a multi-band pass filter (446/532/646, Semrock) and the air objective. Figure 1—figure supplement 2 displays a demo pattern that was projected using our setup, which indicate that the spatial resolution or the contrast of the micro-projection reaches 1.0 μm or 50:1, respectively.

### Manipulation of TFP meditated motility of single cells

Bacterial strain (PAO1-*bphS*-pcdrA-GFP-mCherry) were inoculated into a flow cell (Denmark Technical University) and continuously cultured at 26.0 ± 0.1°C by flowing FAB medium (3.0 mL · h^−1^), in which the flow cell was modified by punching a hole with a 5 mm diameter on the channel, and the hole was sealed by a coverslip that allows the manipulation lights to pass through (Figure 1—figure supplement 2b). The inverted fluorescent microscope equipped with a 100×oil objective and a sCMOS camera was used to collect bright-field or fluorescent images with 0.2 or 1/1800 frame rate respectively. The power density of manipulation lights was determined by measuring the power at outlet of the air objective using a power meter (Newport 842-PE). GFP or mCherry was excited using a 480 nm or 565 nm LED lights (ThorLabs) and imaged using single-band emission filters (Semrock): GFP (520/28 nm) or mCherry (631/36 nm). In our optogenetics manipulation experiments, the motility of single cells were continuously monitored in the absence of optogenetics manipulation using bright-field images in the first hour (Figure 1—figure supplement 1). Real time bacterial tracking algorithm allow us to identify the mobile or immobile TFP meditated motility type. The moving velocity of single cells are directly calculated by |**r**(*t* + Δ*t*) − τ(*t*)|/Δ*t*, where **r**(*t*) is the position of the bacterium at the time *t*, Δ*t* = 30 min. The cells with moving velocity larger than 0.03 μm.min^−1^ were defined to the mobile types. Afterwards, 33-66 % mobile cells were selected to be manipulated using ATI with the illumination at 0.05 mW.cm^−2^, which allowed us to compare the results arising from illuminated or un-illuminated mobile cells in one experiment. The c-di-GMP levels in single cells were gauged using the ratio of GFP and mCherry intensities.

### Guiding biofilm formation using ATI

Bacterial strains of PAO1-*bphS*-EGFP and PAO1-*bphS*-mCherry were 1:1 mixed and inoculated into the modified flow cell to continuously culture at 26.0 ± 0.1°C by flowing FAB medium (3.0 mL · h^−1^). The inverted fluorescent microscope equipped with a 100×oil objective and a sCMOS camera was used to collect bright-field or fluorescent images with 0.2 or 1/1800 frame rate respectively. The cells with green or red fluorescence were selected respectively to be manipulated using ATI with a power density of 0.05 mW.cm^−2^ during the first 10 hours. Afterwards, the flow cell contains the distinctive young biofilms were continuously cultured up to 3 days in the dark to allow these young biofilms to mature. Finally, a laser-scan confocal microscope (Olympus FV1000) equipped with a 100×oil objective was used to image the cells organizations as well as the three-dimensional (3D) structures of the mature biofilms using z-axis scanning (0.5 μm per step). The confocal images acquired in different z-positions were used to reconstruct the structure of mature biofilms using software ImageJ. Experiments of biofilm cultivation were carried out at least three times.

## Acknowledgements

We thank Dr. M. Gomelsky for kindly providing optogenetics plasmid. This work was financially supported by National Natural Science Foundation of China (21474098, 21274141, 21522406) and Fundamental Research Funds for Central Universities (WK2340000066, WK2030020023).

## Competing interests

All authors declare no competing interests.

## Figure Supplements

**Figure 1—figure supplement 1.**
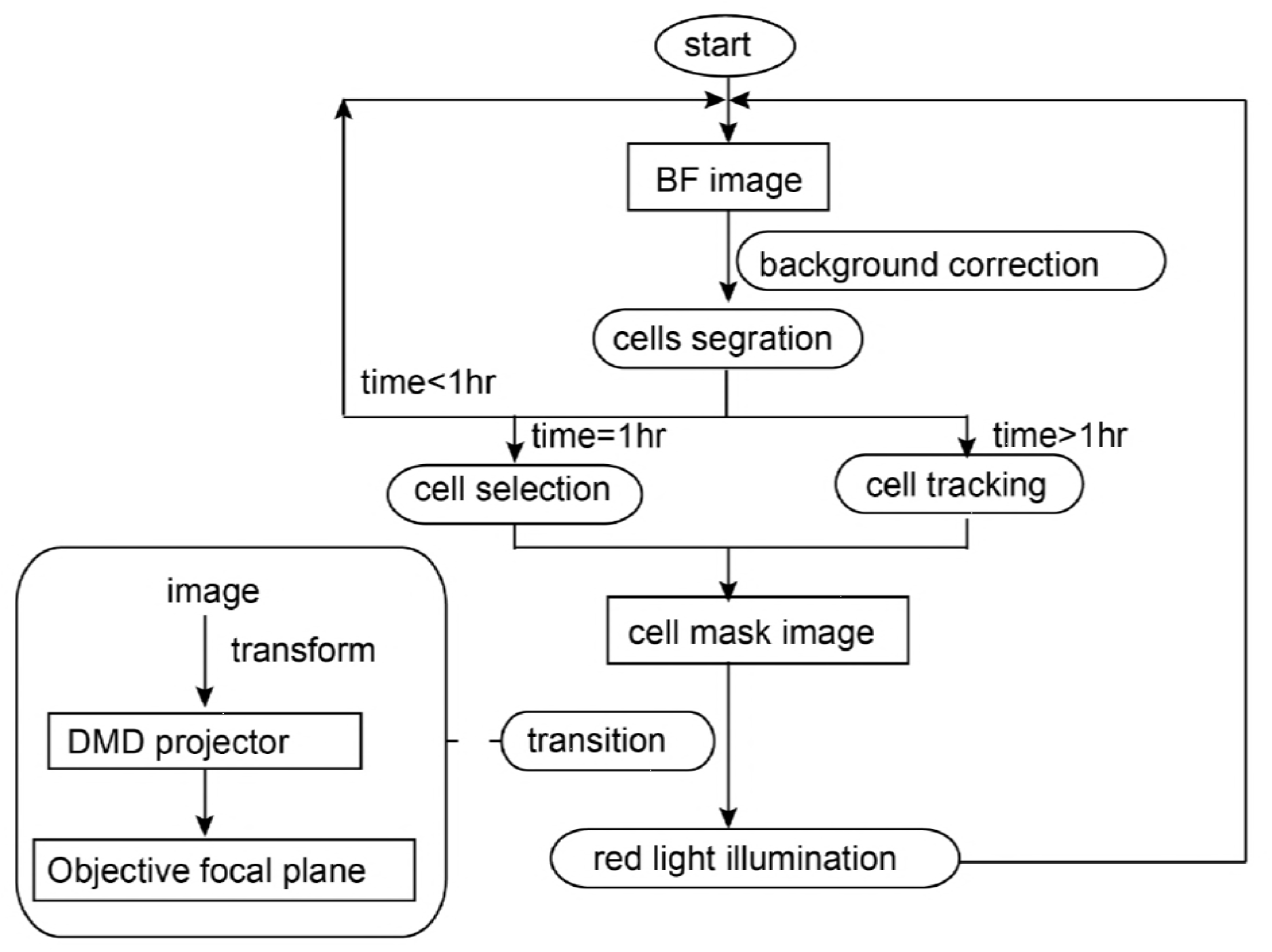
Data analysis with interactive information feedback system. The process was illustrated in a simplified sequence flow diagram. Simply, bright field images were taken to recognize and segment the single bacteria using an image processing algorithm coded by MATLAB; Real time information of the selected cells was secondly transferred to a digital micromirror device (DMD) to generate a mask; The selected cells were finally illuminated using the projected mask through an additional long working distance objective.

**Figure 1—figure supplement 2.**
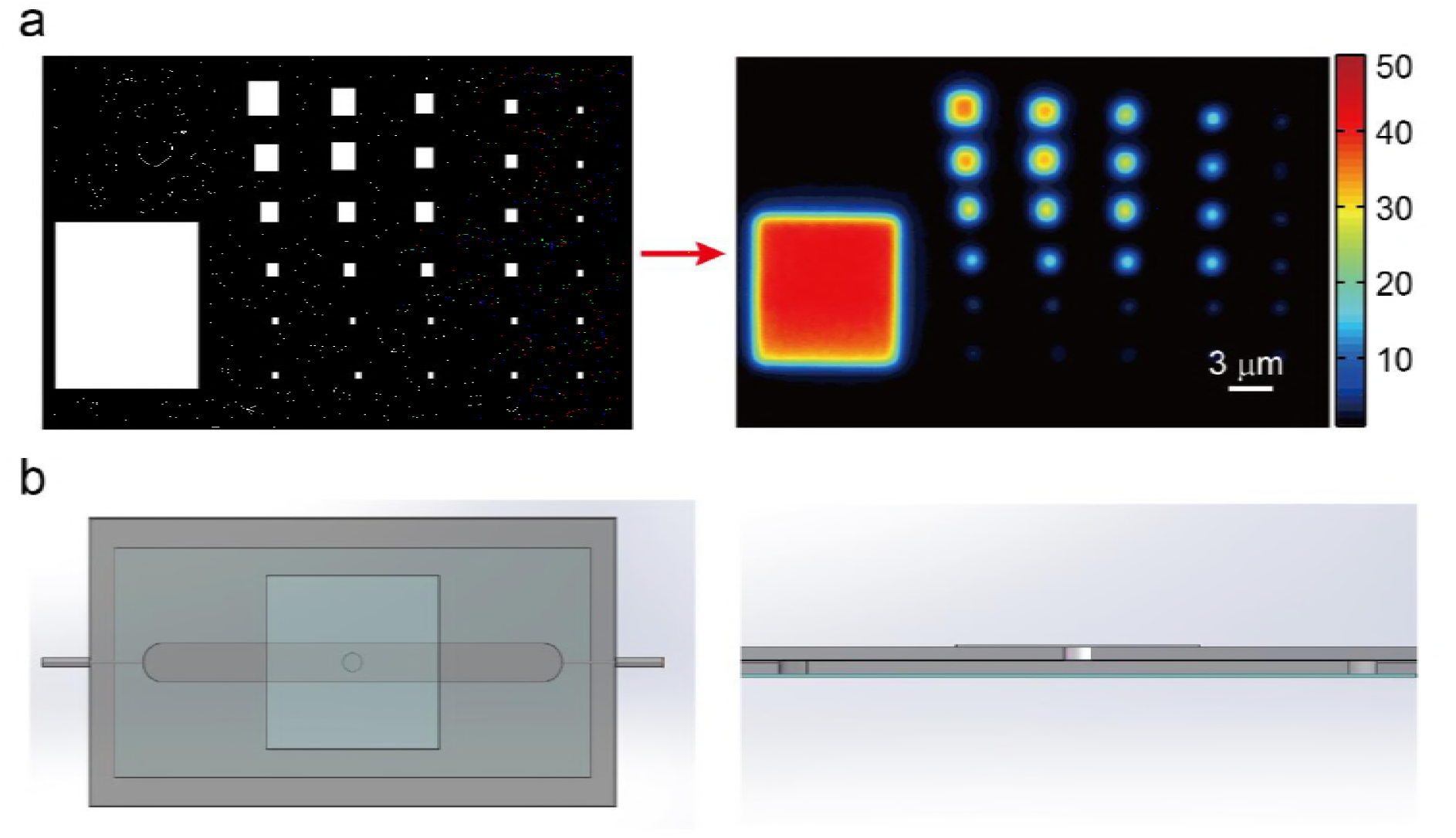
**(a)** A sequence demon patterns with different size were projected using our ATI system. The spatial resolution can reach up to 1.0 μm, which is smaller than length of single *P. aeruginosa* cells (about 2 μm). The ATI system also has high contrast ratio with 50:1. **(b)** Biofilm cultivation was carried out in an adaptive flow cell system. The schematic diagram of the adaptive flow cell is presented.

**Figure 2—figure supplement 1.**
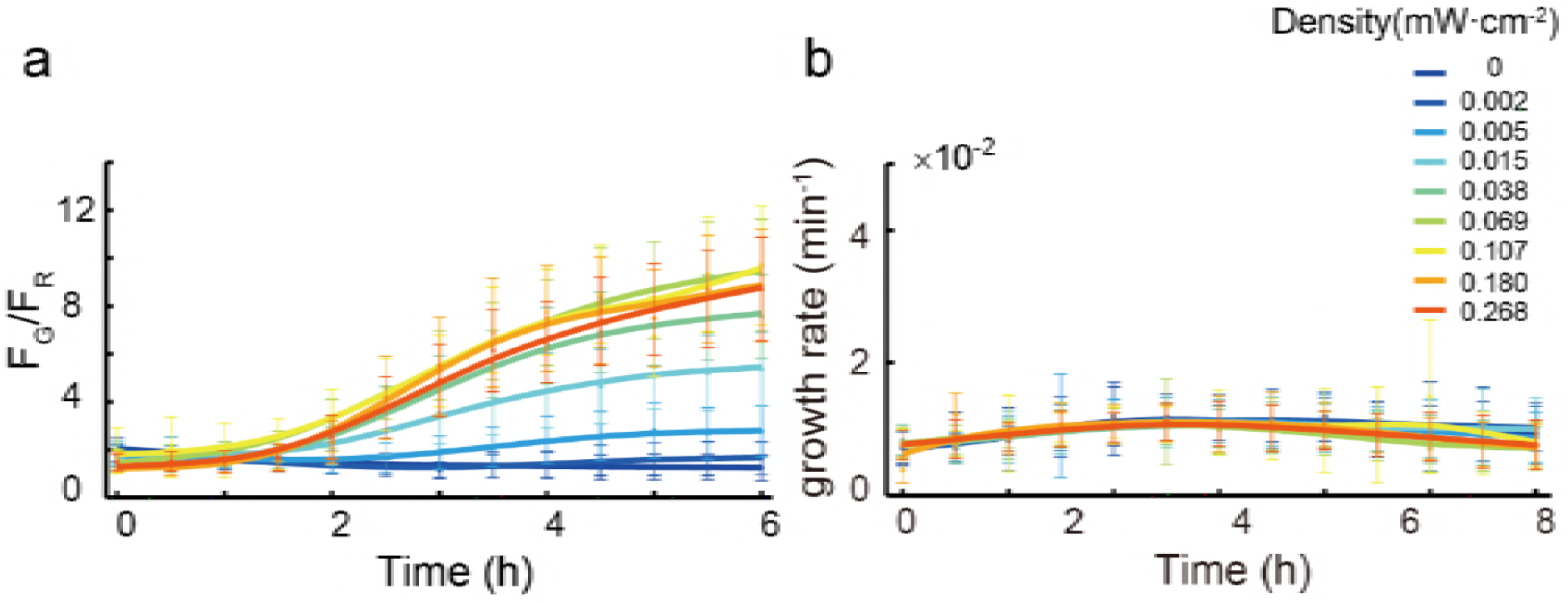
**(a)** Kinetics of Fg/Fr were measured at indicated times with manipulation by ATI under different illumination intensity for 4 hours. **(b)** Relationship between cell growth rate and different illuminations intensity. The intensity being chosen for carrying out experiment (0.05 mW.cm^−2^) does not affect the growth rate of interested cells. Error bars in **a-b** represent means ± s. d with n = 3 biological replicates.

**Figure 2—figure supplement 2.**
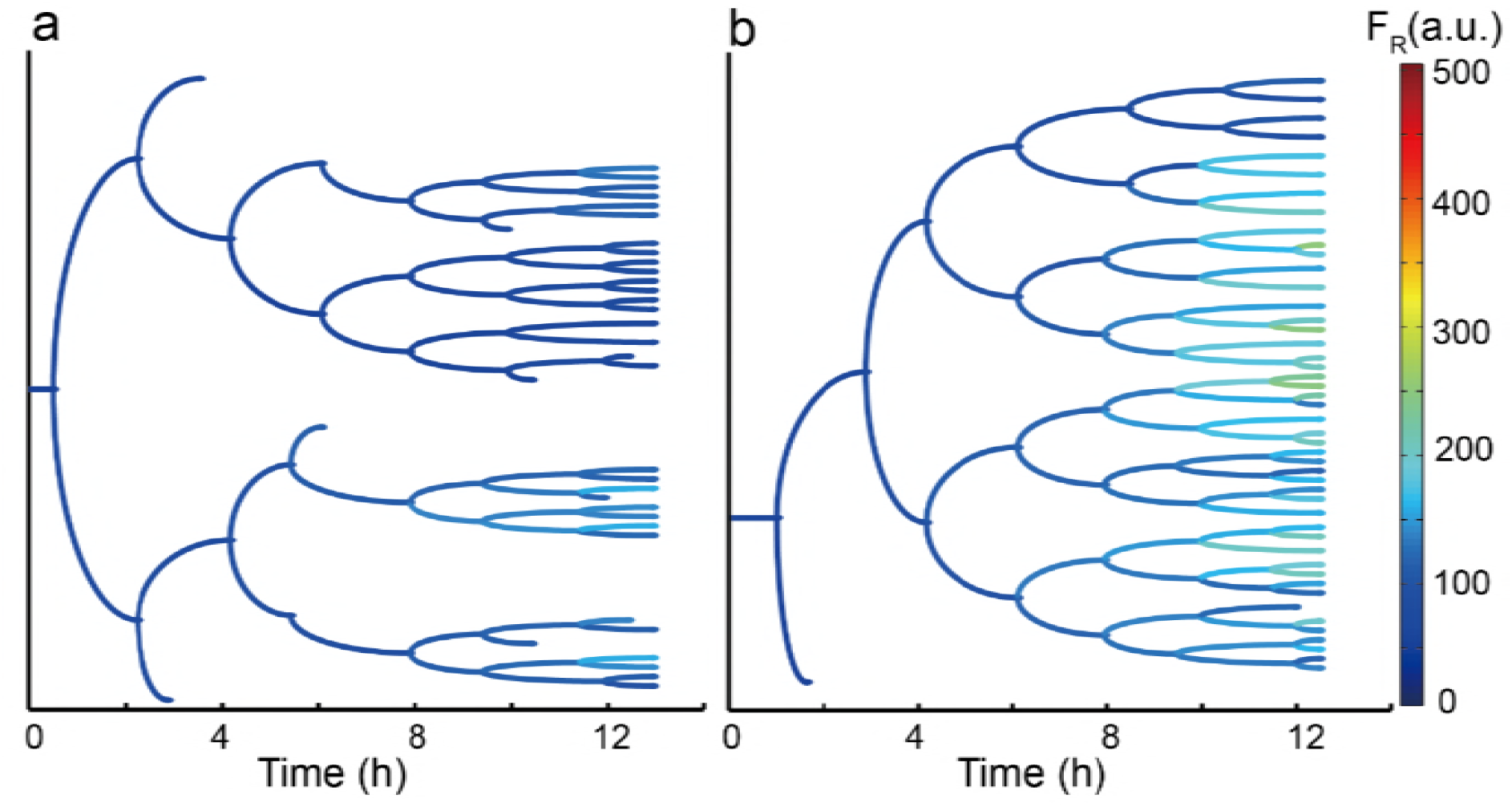
Genealogical tree of one unilluminated cell **(a)** and one illuminated cell **(b)** was used to display the mCherry fluorescence intensity of its offspring separately. The fluorescence arose from mCherry, which is used as an internal control, nearly remains consistent in daughter cells with illuminated or unilluminated.

**Figure 4—figure supplement 1.**
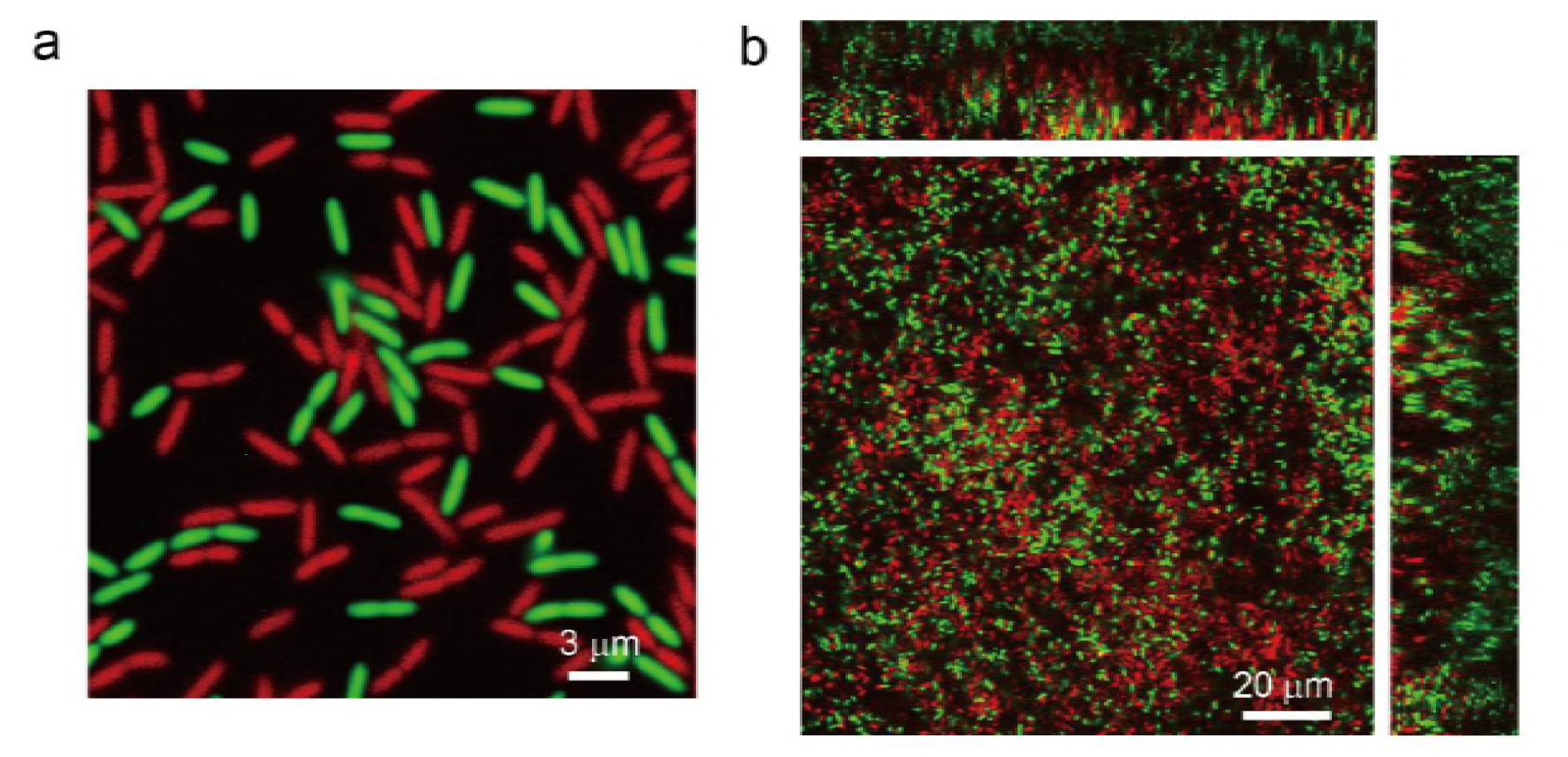
Two different labeled cells with mCherry and EGFP were cocultured in a flow cell to allow them to form biofilms. No cells are selected to be manipulated by ATI. **(a)** and **(b)** were captured at t~10 h or 3 days, respectively.

## Video Legends

**Video 1**. One interested cell being tracked and projected in real time is depicted. Red color represents the region of red LED illumination.

**Video 2**. One interested cell being tracked and projected in real time is depicted. Red color represents the region of red LED illumination.

**Video 3**. Single cells are precisely illuminated by ATI via *in situ* analyzing and tracking bacteria. The left panel shows the merged images of GFP mut3* and mCherry fluorescence microscopy images changed over time. The right panel shows the merged images of red LED projected patterns and bright field images as the same to left panel. The fluorescence intensity of GFP mut3* in those illuminated cells and their offspring (as shown red color merged in right panel) is significantly increased after using ATI for 7 hours, which is sharply contrast to that the fluorescence intensity of GFP mut3* in those unilluminated mobile cells remain in low.

